# A Drug-free, Self-destruction Strategy to Combat Bacterial Infections by Using the Intrinsically formed Magnetic Nanoparticles in Bacterial Pathogens

**DOI:** 10.1101/2021.02.03.429514

**Authors:** Swati Kaushik, Jijo Thomas, Vineeta Panwar, Preethi Murugesan, Vianni Chopra, Navita Salaria, Rupali Singh, Himadri Shekar Roy, Rajesh Kumar, Vikas Gautam, Deepa Ghosh

## Abstract

The growing number of multiple drug resistant (MDR) bacteria and the dwindling pipeline of new antibiotics are driving us towards a ‘post-antibiotic era’ in which even common infections would become difficult to treat. To address this, an antibiotic-free strategy that can combat multiple bacteria is recommended. Most of the proposed approaches nevertheless have several limitations, including bacterial targeting. To overcome such limitations, the proposed strategy employs the bacterial machinery to self-destruct. Herein, the biosynthesis of magnetic nanoparticle (MNP) is reported for the first time in multiple pathogenic bacteria, including MDR bacteria. The intracellular MNPs composed of superparamagnetic zinc ferrites were formed in presence of iron and zinc precursors. Exposure of the treated bacteria/biofilms to an alternating magnetic field (AMF) exhibited hyperthermia (5-6°C) and a dramatic decrease in bacterial viability, suggesting the MNPs therapeutic potential. Likewise, the bacteria existing *in vivo* biosynthesize the MNPs by mining these elements from the host. To determine its therapeutic efficacy, the infected tissues were exposed directly to AMF. A 3-4 log reduction in bacterial burden, as compared to antibiotics treatment, confirmed the significance of using naturally existing MNPs to combat bacterial infections. The proposed broad–spectrum approach can therefore aid in overcoming the challenges facing anti-bacterial therapies.

## 1. Introduction

The ongoing pandemic has so far affected 175,987,176 and resulted in 3,811,561 deaths ^[1]^. With viral respiratory infections known to make the host more susceptible to microbial infections ^[2]^, the reports on COVID-19 patients provide ample evidence of the wide spread use of antibiotics to control the secondary bacterial infections ^[3]^. The most frequently detected causative agents in such infections generally belong to a class of bacteria that are responsible for nosocomial infections (*Staphylococcus* spp., *Escherichia coli, Pseudomonas* spp., *Enterococcus* spp., *Klebsiella pneumoniae, Acinetobacter* spp., *Enterobacter* spp., etc) in hospitalized patients ^[3d, 4]^. Many of these strains are resistant to most antibiotics ^[5]^. The widespread use of antimicrobial soaps, disinfectant cleaners and antibiotics during the pandemic is expected to amplify the emergence of additional strains of resistant pathogens that would further impact the existing overburdened healthcare systems ^[6]^. Thus, apart from the projected MDR pathogens contributing to 10 million deaths by 2050 (WHO)^[7]^, the ongoing pandemic is expected to contribute to an exponential increase in the number of fatalities and possibly fuel the next epidemic.

Despite best practices to control the indiscriminate use of antibiotics and antibiotic stewardship, the emergence of new resistant strains and the depleting pipeline of new antibiotics continue to worsen the outlook for anti-bacterial therapies. Various new interventions are being assessed to address the resistant strains ^[8]^. Among the nanotechnology-based anti-microbial approaches, the use of nanoparticles appears to be the most promising ^[9]^. Nanoparticles have been widely explored to improve drug uptake, destroy biofilm formation and target drugs to the site of infection ^[10]^. The use of metal nanoparticles for photodynamic and photo thermal therapies are emerging as complementary strategies to antibiotics, to combat/overcome bacterial resistance ^[11]^. Whereas most of these approaches are aimed to prevent resistance development ^[9] [10b]^, they might be unable to address the bacteria that are already multi drug resistant (MDR). Hence, it would be beneficial to develop an anti-bacterial strategy that can counter a wide spectrum of both sensitive and resistant bacterial strains.

Most bacteria in the body thrive between 33-41°C. Elevated temperatures impede bacterial proliferation through autolysis and cell wall damage ^[12]^. Hyperthermia, the process of heating tissues to a high temperature (45-50°C) for a short duration, is widely used in clinical practice. Among the various proposed hyperthermia techniques, heat generation using magnetic nanoparticles (MNPs) is considered safe as it absorbs electromagnetic radiation and converts it to localized heat on exposure to an alternating magnetic field (AMF). As the MNPs impart heat closer to cell membranes, a mild-moderate heating is adequate to induce significant cell death ^[13]^. Most MNPs that were employed in anti-bacterial studies were prepared using a synthetic route ^[14]^. The major hurdles facing such MNPs include its aggregation, heating efficiency, bacterial targeting, apart from toxicity when used in large scale ^[15]^. Despite the promise of antibody-functionalized MNPs to target the bacteria, the approach is bacteria-specific and expensive ^[14a]^. Unlike synthetic MNPs, the monodispersed biosynthesized MNPs have better heating efficiency due to their narrow size range ^[12, 16]^. In nature, the biosynthesized MNPs are present inside the bacteria only in a special class of bacteria termed, Magnetotactic bacteria ^[17]^. These fastidious bacteria have an extremely slow rate of bio mineralization (>1 week) ^[18]^. Apart from magnetotactic bacteria, another class of bacteria termed ‘iron reducing bacteria’ biosynthesize MNPs extracellular in presence of iron precursors ^[19]^. Despite the efficacy of these biosynthesized MNPs to kill bacteria ^[20]^, its clinical application is hindered by challenges that are similar to synthesized MNPs.

The biosynthesized MNPs in magnetotactic bacteria are generally made of oxides of iron including magnetite, maghemite or gregite ^[21]^. A recent report demonstrated the formation of extracellular zinc ferrites (ZnFe_2_O_4_) by an iron reducing bacteria, *Geobacter sulfurreducens* in presence of iron and zinc ^[22]^. With an objective of developing a strategy that overcomes the limitations of existing MNPs to combat bacteria, we aimed to induce pathogenic bacteria to biosynthesize MNPs. Here we present evidence of intracellular MNP formation in nosocomial infection causing bacteria like *S. aureus, E. coli, P. aeruginosa* and *K. pneumoniae* on treatment with iron and zinc precursors. The MNPs in the bacteria were characterized using multiple techniques. The superparamagnetic nature of the MNPs was confirmed using Superconducting Quantum Interference Device (SQUID). Magnetotaxis of the nanoparticles and bacteria was visualized using bright-field microscope. The hyperthermia response of the MNPs was studied using alternating magnetic field (AMF). The host defense system in the body prevents bacterial access to these elements through a process termed nutritional immunity ^[23]^. Nevertheless, to facilitate their survival and replication, the infectious pathogens have developed smart strategies to mine these metals from the host ^[24]^. In this report we present evidence of virulent bacteria in the host to similarly biosynthesize MNPs with the acquired iron. The enhanced susceptibility of these bacteria to magnetic hyperthermia, in comparison to antibiotics, suggested the therapeutic potential of the naturally existing MNPs to induce efficient hyperthermia leading to bacterial death.

## 2. Results and Discussion

Bacteria have evolved multiple strategies to acquire and regulate iron and zinc to build a virulence arsenal that can challenge even the most sophisticated immune responses ^[23]^. As both these elements are essential for bacterial virulence, we evaluated the response of bacteria (*S. aureus, E. coli, P. aeruginosa and K. Pneumoniae*) to a broth containing FeCl_2_ and zinc gluconate. In comparison to the untreated control and treatment with either Fe/Zn, the cultures containing both Fe and Zn developed a dark brown color, suggesting iron oxide formation **(Figure S1)**. The presence of iron oxide in the bacteria was confirmed using Perl’s Prussian blue staining ^[25]^**(Figure. S2)**. The response of the intracellular iron oxide to magnetic field was evaluated by exposing the bacterial lysates to a magnet. The alignment of the particles along the magnetic field lines indicated the presence of magnetic particles in the treated bacteria **(Figure S3)**. Magnetic field-induced motility was observed in the bacteria treated with Fe & Zn **(Video M2, M4, and M6)** as compared to untreated control **(Video M1, M3 andM5)**. The motility was similar to the magnetotaxis reported in magnetotactic bacteria ^[27]^. IC-PMS was used to assess the intracellular Fe and Zn content in the bacteria. As seen in table ST1, the microbes treated with both iron and zinc exhibit higher Fe content, indicating the influence of Zn on Fe uptake **(Figure S3)**.

Further characterization was performed using a representative Gram-positive (*S. aureus*) and Gram-negative bacterium (*E. coli*). TEM analysis of Fe and Zn treated *S. aureus* and *E. coli* revealed the presence of a large number of nanocrystals distributed throughout the respective bacteria **(Figure 1 B & F)**, while no such phenomenon was seen in untreated control **(Figure 1 A & E)**. On further analysis of the treated bacterial lysates, Quasi-cuboidal shaped particles **(Figure 1I)** of 13-19 nm size were observed **(Figure 1J)** ^[26]^. HR-TEM images of individual nanocrystals showed the lattice fringes, indicating the crystalline nature of the material ^[27]^. The inter-planar spacing calculated from these fringes exhibited d-spacing of 0.283 nm and 0.489 nm corresponding to 100 planes of ZnO and 111 planes of zinc ferrite (ZnFe_2_O_4_) respectively ^[26, 28]^ **(Figure 1C&G)**. The diffraction spots indexed from selected area electron diffraction pattern (SAED) confirmed its crystalline nature having (111), (311), (220), (400), (511), (440) planes of ZnFe_2_O_4_ and (100), (101), (110) plane of ZnO ^[29]^**(Figure 1 D & H)**. EDAX analysis **(Figure 1K)** and elemental mapping **(Figure S4)** confirmed the presence of iron, zinc and oxygen with homogeneous distribution of the above-mentioned elements.

**Figure 1:**
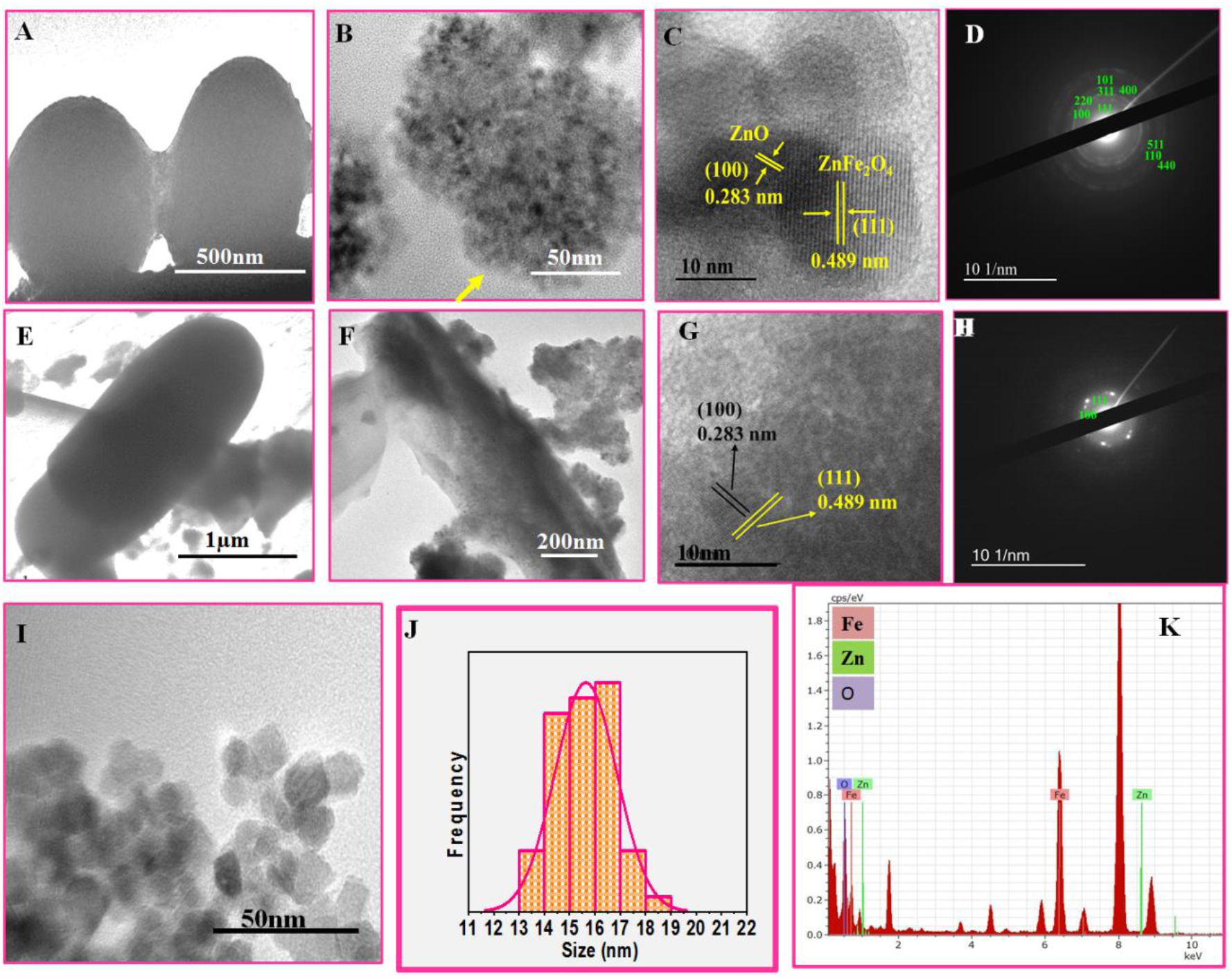
Characterization of nanoparticles. The upper and lower panel represent TEM images of *S. aureus* and *E. coli* respectively. (A, E) Untreated bacteria; (B, F) Bacteria treated with FeCl2 and zinc gluconate (arrow indicates nanoparticles); (C, G) HR-TEM showing D-Planar spacing of nanoparticles; (D, H) SAED pattern of nanoparticles; (I) TEM image of nanoparticles; (J) Size distribution of the nanoparticles (K) EDAX showing the presence of Fe, Zn and O obtained from *S. aureus*.

To evaluate the magnetic nature of the nanoparticles, field dependent magnetization measurements were performed on the lyophilized samples of treated bacteria using SQUID. Figure 2 displays the magnetic properties (M-H loop at 300 K (A), at 5 K (B) and M-T curve (C) with *S. aureus*. At room temperature (300 K), negligible coercivity was observed signifying the superparamagnetic behavior of the nanoparticles due to zero coercivity **(inset, Figure 2A)**. The magnetic signal observed at 5K displayed low coercivity (150 Oe**) (inset Figure 2B)**, confirming the characteristic properties of superparamagnetism ^[30]^. The zero field cooled (ZFC) and field cooled (FC) curves of the samples obtained at an applied field of 500 Oe further confirmed the superparamagnetic behavior with blocking temperature at ∼ 150 K **(Figure 2C)**. A similar observation was observed with *E. coli* **(Figure S5)**. As the superparamagnetism characteristics are generally exhibited by MNPs that exist as single domain particles, it can be concluded that the magnetic particles present in the treated bacteria are also single-domain superparamagnetic nanoparticles ^[31]^. Additionally, zinc ferrite nanoparticles are reported to have better magnetization properties than iron oxide nanoparticles, which could be responsible for the observed magnetotaxis ^[22]^.

**Figure 2:**
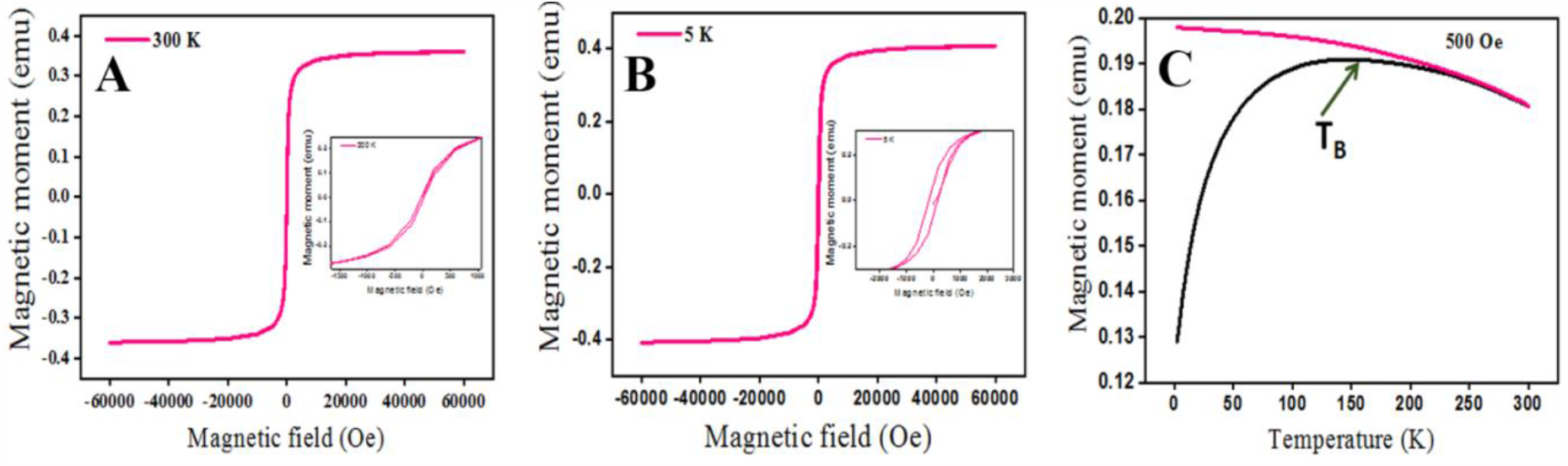
Magnetic measurement of nanoparticles in *S. aureus*. FeCl_2_ and zinc gluconate treated bacteria were lyophilized and magnetization versus magnetic field was measured at (A) 300 K, (B) 5 K and inset represents coercivity (C) Measurement of temperature dependence of magnetization (FC/ZFC curves).

**Figure 3:**
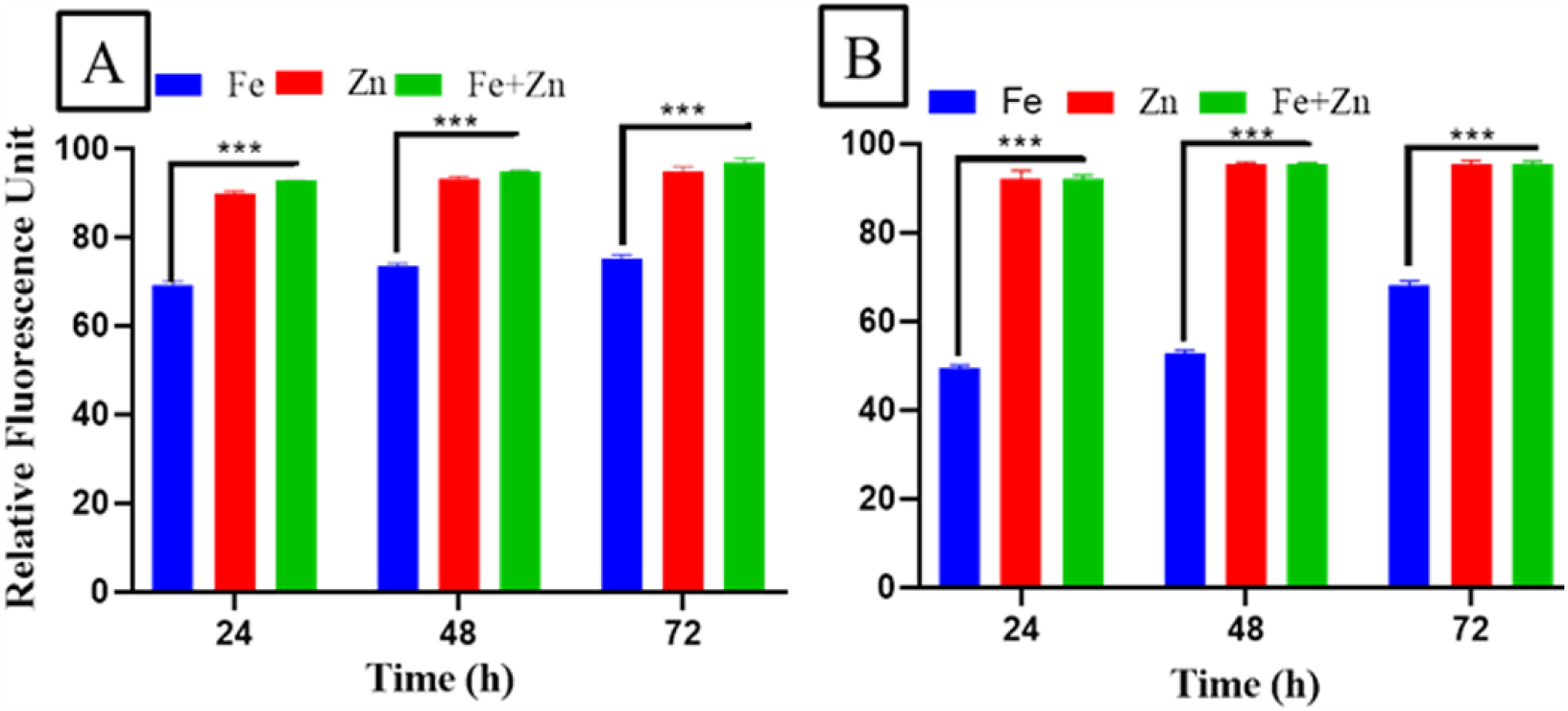
Time dependent ROS expression. (A) *S. aureus* and (B) *E. coli* treated with FeCl2 /zinc gluconate/both for 24-72 h. The data represents the mean ± SD obtained from 3 experiments in which N=3 in each group. Two-way ANOVA Bonferroni multiple comparison test where *** represents p-value <0.001.

XRD analysis of the treated bacteria revealed the presence of crystalline matter **(Figure S6A)**. The numerous strong Bragg reflections could be indexed to both ZnO and zinc ferrite phases^[28]^ respectively, supporting the TEM results. The crystalline peaks at 2θ = 31.73°, 36.2° and 56.6° represents (100), (101), (110) hkl plane correspond to the hexagonal crystal structure of ZnO according to JCPDS no. 36-1451 ^[32]^. The peaks at 2θ = 18.3°, 30.1°, 35.2°, 43.1°, 53.3°, 56.7° and 62.4° with hkl plane of (111), (220), (311), (400), (422), (511) and (440) correspond to the cubic spinel structure of ZnFe_2_O_4_ according to JCPDS no.79-1150 ^[29]^. Fourier transform infrared (FT-IR) analysis of the lyophilized treated-bacteria also exhibited peaks in the region from 670-550 cm^-1^ reflecting the stretching vibration mode associated to Fe–O bonds in the crystalline lattice of ZnFe_2_O_4_ ^[27]^. The other peaks observed in the region from 476-417 cm^-1^ indicated the presence of Zn-O bond **(Figure S6 C&D)**. The presence of N-H stretch between 3,500-3,100 cm^-1^ in the full scan FTIR spectra could be contributed by the proteins present in the bacterial cells **(Figure S6B)**. These results reconfirmed the presence of zinc ferrite and zinc oxide in the treated bacteria.

Iron uptake in the bacteria triggers the Fenton/Haber-Weiss reaction resulting in the formation of superoxide, hydrogen peroxide and hydroxyl radical ^[33]^. In presence of these reactive species (ROS), the soluble Fe^2+^ is converted to insoluble oxides of Fe^3+ [34]^. We evaluated ROS levels in bacteria treated with FeCl_2_ / zinc gluconate and their combination. In comparison to iron alone, the bacteria treated with zinc, and in combination with iron, displayed a substantial increase in the ROS levels, suggesting zinc as a potent ROS inducer **(Figure3)**. As ROS is known to directly influence biomineralization, an increase in ROS observed in Fe and Zn-treated bacteria could be responsible for MNP formation. This data is supported by a recent report showing Zn to increase both intracellular Fe and oxidative stress in *E. coli* ^[18]^. Additionally, we observed microbes expressing increasing levels of ROS to contain higher intracellular iron accumulation **(Table ST1)**. In a recent study with magnetotactic bacteria (*Burkholderia* sp.), Pan *et al* had concluded that the MNPs in the bacteria help in the efficient scavenging of excess ROS, by the antioxidase-like activity of iron crystals ^[35]^. The MNPs observed in the pathogenic bacteria might also follow a similar strategy to overcome ROS-induced toxicity.

To date, magnetotactic bacteria (MTB) are the only naturally occurring bacteria that are known to form intracellular MNPs and exhibit magnetotaxis in response to magnetic field ^[36]^. MNP containing magnetosomes-derived from MTB are known to induce magnetic field-dependent temperature rise on exposure to AMF ^[37]^. MNPs with diameter <15 nm is considered most effective for generating hyperthermia ^[38]^, and in comparison to magnetite, zinc ferrite nanoparticles have shown improved magnetization properties ^[22]^. As the biosynthesized MNPs are made of zinc ferrite in the size range of 13-19 nm, we evaluated its heating potential. Exposure of the treated bacteria to AMF at two different frequencies i.e. 164 KHz and 347 KHz for 30 min resulted in a time dependent increase in media temperature, with an improved heating outcome of ∼6°C at 347 KHz **(Figure 4A)**. To check the bacterial susceptibility to hyperthermia, the bacterial suspension before and after the respective AMF treatment was spread on agar plates and the colonies checked after 24 h **(Figure 4B & 4C)**. The significant temperature rise observed with the higher frequency, translated to increased cell death as compared to lower frequency. The data indicated that the biosynthesized MNPs were efficient to cause magnetic-hyperthermia induced cell death **(Figure 4D)**. Field emission-scanning electron microscopic images revealed the distorted morphology of the bacteria post-AMF **(Figure 4Eb)** as compared to pre-AMF **(Figure 4Ea)**. Among the two strains that were tested, *S. aureus* (ATCC 25923) is reported to be sensitive to vancomycin and ciprofloxacin (MIC 1-2 μg/ml), while *E. coli* (ATCC 25922) is resistant to vancomycin (MIC >64 μg/ml) but sensitive to ciprofloxancin (MIC 0.008-1 μg/ml) ^[39]^. On evaluation of the bacterial susceptibility to vancomycin and ciprofloxacin, we observed similar results (data not shown). The reduction in cell numbers observed with AMF in the above bacteria implied, the proposed approach can be effective against both Gram-positive and Gram-negative bacteria; as well as in drug sensitive as well as drug resistant bacteria.

**Figure 4:**
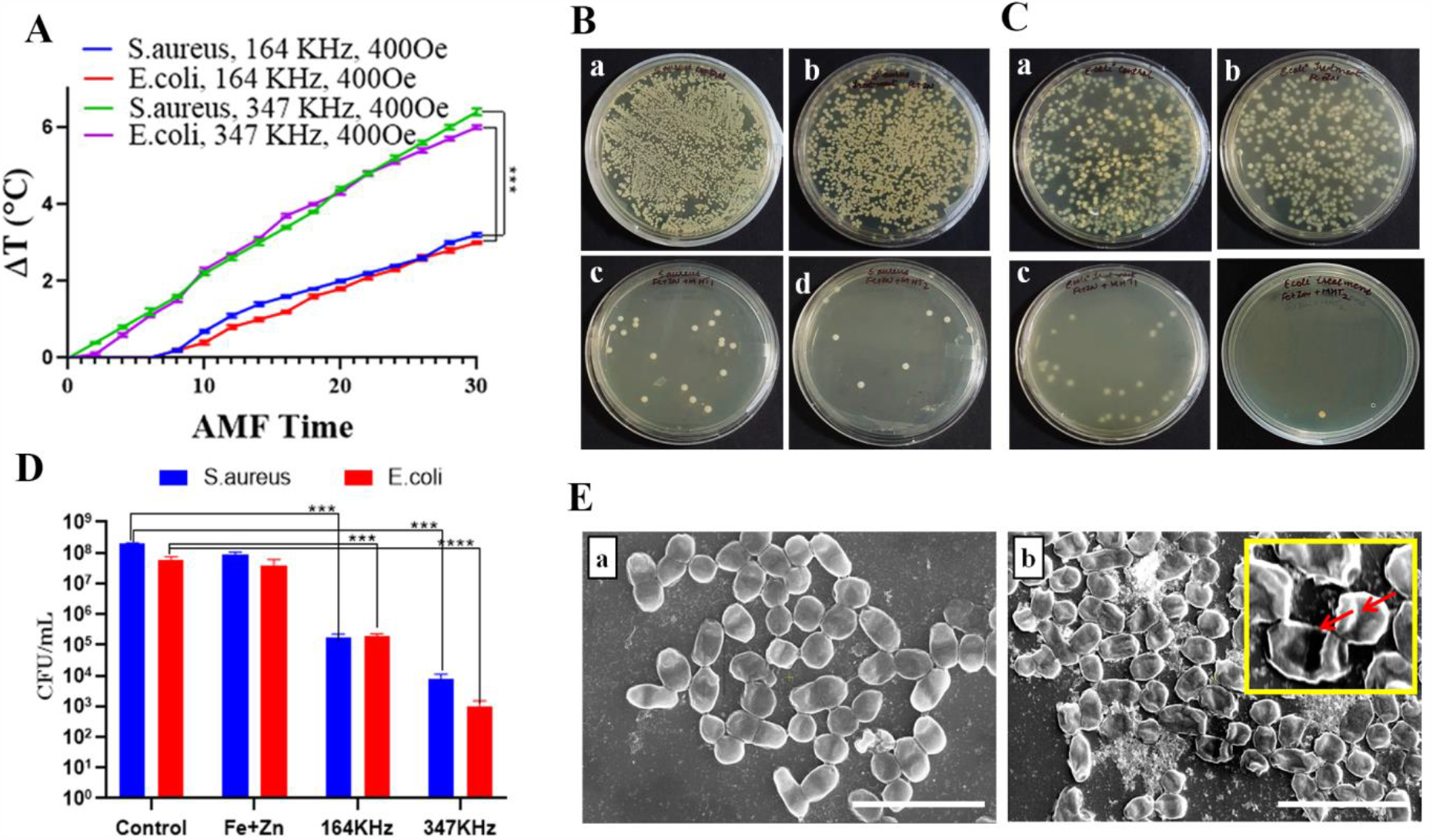
Bacterial response to hyperthermia. (A) Time and magnetic-field-dependent temperature rise in *S. aureus* and *E*.*coli*. Bacterial cell colonies obtained with respective treatments. (B) *S. aureus* and (C) *E. coli*, wherein, (a) represents untreated bacteria (b) bacteria treated with FeCl2 and zinc gluconate (c) Treated bacteria exposed to AMF at164 KHz, 400 Oe and (d) Treated bacteria exposed to AMF at 347 KHz, 400 Oe; (D) Number of bacterial colonies obtained after respective AMF treatment. The data represents the mean ± SD obtained from 3 experiments in which N=3 in each group. Two-way ANOVA Bonferroni multiple comparison test where *** represents p-value <0.001 and **** represents p-value<0.0001. (E) FE-SEM images of treated bacteria (a) before and (b) after AMF. Scale bar represent 5µm.

Biofilms are resistant to antibiotics and host immune responses due to the impervious protective material laid down by the surface adhered bacteria ^[40]^. As these are often associated with persistent infections, several nanotechnology based approaches have been attempted to prevent/destroy biofilms ^[25b, 41].^ Hyperthermia using MNPs that have been modified to facilitate its penetration of biofilms have been employed to destroy biofilms ^[42]^. To evaluate the MNP formation in biofilm, we prepared biofilms using *S. aureus* and *E. coli* and treated with Fe and Zn precursors for 36 h. The treated biofilms were exposed to AMF for 30 min and the viability was evaluated using live/dead staining with FDA/PI staining. The absence of green cells (FDA) in AMF exposed biofilms **(Figure 5A&B)**, indicated the complete destruction of the bacteria in the biofilms. The susceptibility of the bacteria in the biofilms is due to MNP – induced magnetic-hyperthermia and establishes the potential of using the above strategy to effectively destroy biofilms.

**Figure 5:**
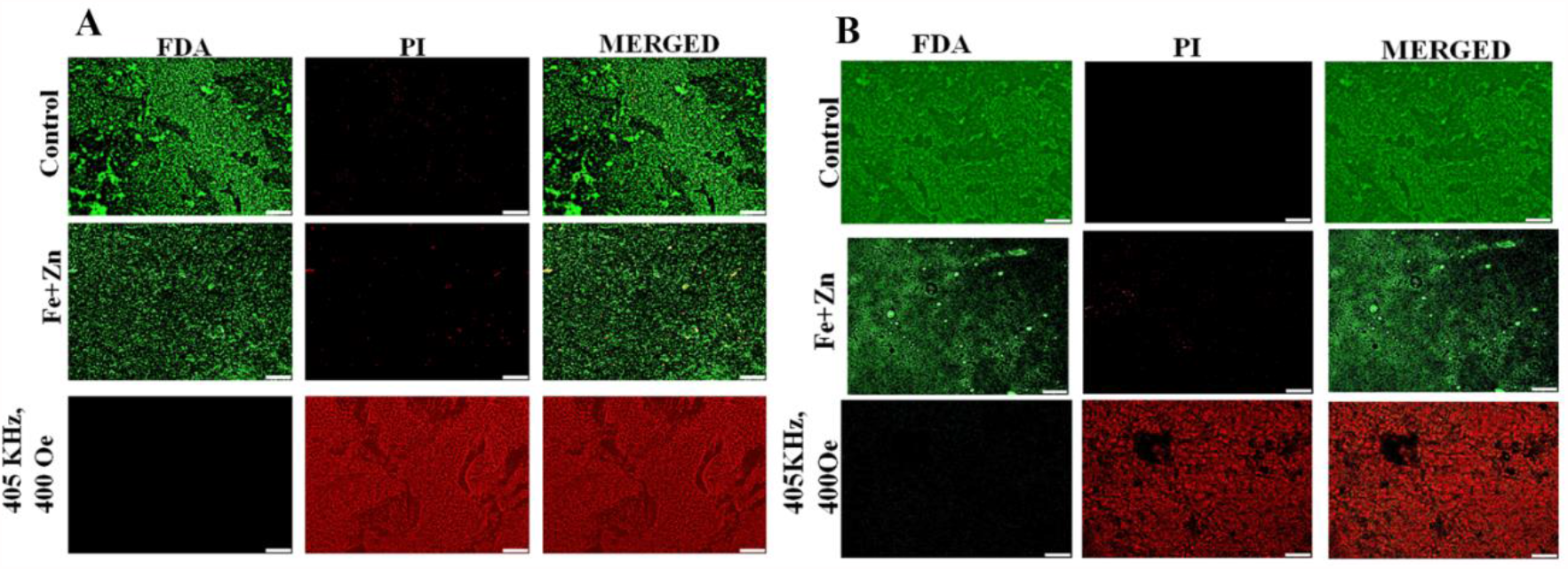
Effect on biofilms. Biofilms of (A) *S. aureus* (B) *E. coli* bacteria were treated with Fe and Zn precursors for 36 h and exposed to AMF for 30 min. The biofilms were treated with FDA/PI and the live (green) /dead (red) cells were visualized using fluorescence microscope. Scale bar represent 100 µm.

Virulent bacteria overcome the host’s nutritional immunity and gain access to elements like iron and zinc. Bacterial uptake of these elements is regulated by the metal-dependent Fur/Zur family of proteins which control the respective genes involved in their acquisition. Besides normal physiological processes, these regulatory proteins are also responsible for the expression of virulence factors ^[43]^. To confirm if the bacteria in the host harbor similar MNPs, we examined the bacteria from various infected human specimens **(Table ST2)** for the presence of iron oxide. The pathogenic bacteria in each specimen was identified by Gram-staining and stained with Perl’s Prussian blue **(Figure S7)**. A blue color in the bacteria (*E*.*coli, S. aureus, A. baumannii, K. pneumoniae, S. epiderdimis, S. hemolyticus etc*.) signified the existence of iron oxide, an observation that was similarly noted in the bacteria treated with Fe & Zn **(Figure S2)**. Notable is that most of these bacteria are listed by the Center for Disease Control (CDC) as nosocomial pathogens related to Healthcare-associated infections. Many of these are resistant to multiple drugs resulting in high patient morbidity and mortality ^[44] [60]^. TEM analysis of the infected samples revealed several nanocrystals distributed in the bacteria **(Figure 6Aa)**. Analysis of the carbonized specimens showed spherical crystalline nanoparticles of iron oxide ∼ 20 nm **(Figure 6Ab)**. The D-planar spacing and the crystalline structure **(Figure 6Ac & 6Ad)** confirmed the presence of iron NPs. This is the first report that demonstrates the natural existence of iron NPs in bacteria thriving *in vivo*. The magnetic property of the iron nanoparticles was confirmed from the microscopic evidence of nanoparticle aggregation observed in the lysate on exposure to magnet **(Figure S8)**. To determine if the MNPs present in the virulent bacteria can generate heat with AMF, we exposed the RBC-depleted infected blood samples to AMF at 347 KHz, 400 Oe for 30 min. A time-dependent rise in temperature observed in all the samples, confirmed the magnetic hyperthermia potential of the MNPs **(Figure 6B)**. Despite the difference in the temperature rise observed between various strains, a 2-4 log reduction in the bacterial colonies was observed **(Figure 6D)**. The bacterial colonies obtained before and after AMF exposure are displayed in **(Figure 6C)**. The susceptibility of the above bacteria to antibiotics was evaluated using vacomycin and ciprofloxacin respectively (2 µg/mL). While *S. epiderdimis*, responded only to ciprofloxacin, it was resistant to vancomycin. The other bacteria were resistant to both the antibiotics **(Figure S9)**. Considering the significantly higher reduction in bioburden with AMF as compared to antibiotics treatment, it can be concluded that AMF alone can play an important role in combating bacterial infections.

**Figure 6:**
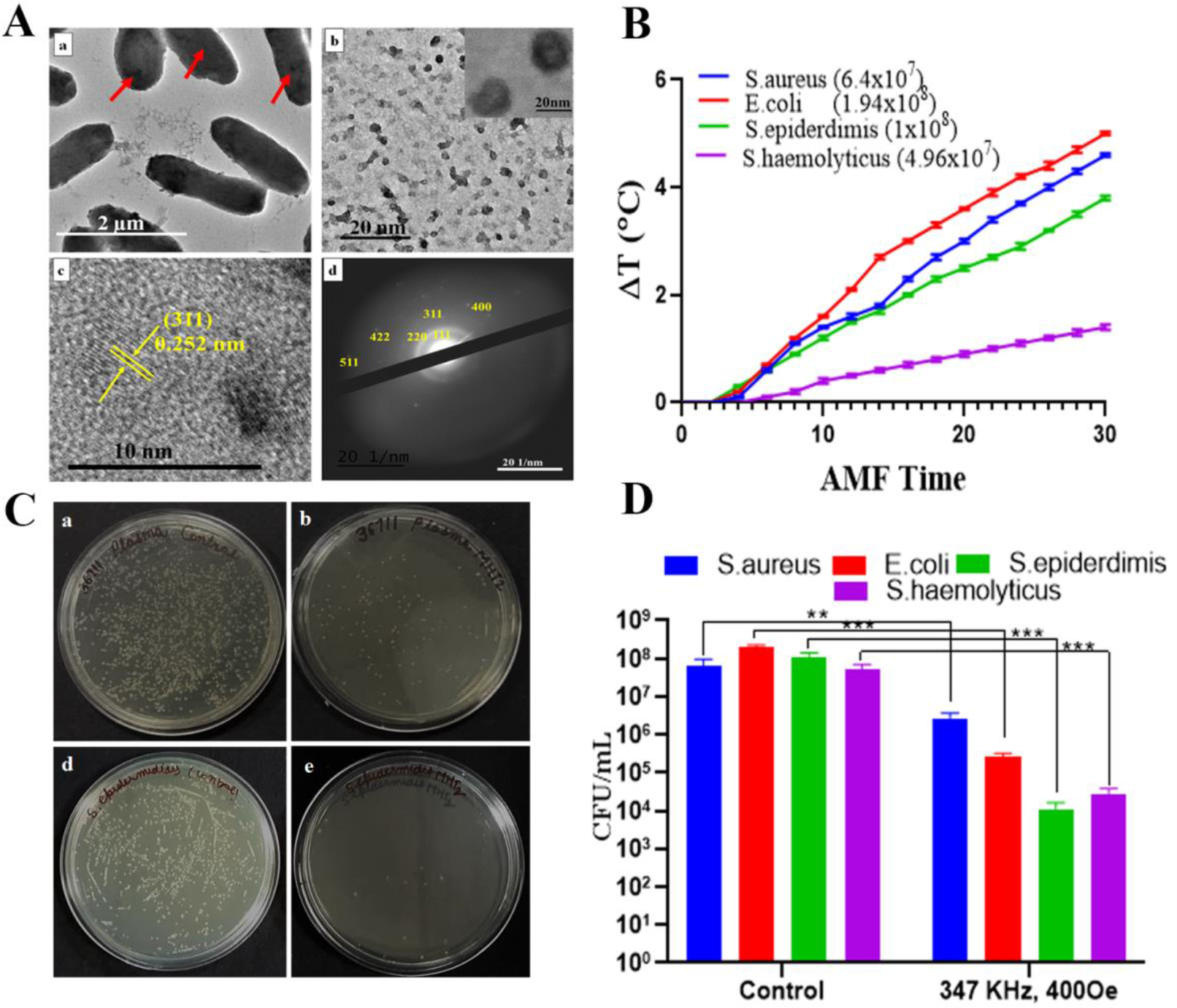
Characterization and hyperthermia response of bacteria derived from infected human samples. (A) Bacteria isolated from blood specimen was characterized using TEM. (a) *E. coli* containing nanoparticles (indicated by red arrows); (b) Purified nanoparticles obtained from the above *E*.*coli* sample (inset shows magnified nanoparticles); (c,) HR-TEM showing D-Planar spacing of the nanoparticles; (d) SAED pattern of nanoparticles. (B) Time and magnetic-field-dependent temperature rise in the respective bacteria. The number of respective bacteria is mentioned in brackets. (C) Bacterial colonies obtained post-AMF where, upper panel shows *S. aureus* and lower panel is *S. epiderdimis*, wherein, (a, c) represents untreated bacteria (b, d) bacteria exposed to AMF at 347 KHz, 400 Oe; (D) Number of bacterial colonies obtained after AMF treatment. The data represents the mean ± SD obtained from 3 experiments in which N=3 in each group. Two-way ANOVA Bonferroni multiple comparison test where ** represents p-value <0.01 and *** represents p-value <0.001.

To assess whether AMF can reduce the bacterial load of infected solid tissues, the specimens were treated directly with AMF (347 KHz, 400 Oe) for 30 min. The bacterial load of pre-and post-AMF exposed infected tissues (abdominal muscle and bone specimens) was evaluated after tissue homogenization. A considerable reduction in the bacterial colonies was noted in the AMF-treated tissues, suggesting the penetration of AMF to induce cell death in bacteria located in deep tissues **(Figure S10 C&D)**. Since all virulent bacteria assimilate iron and zinc, it is proposed that intracellular MNPs might exist in all such bacteria. Given the promising anti-bacterial effect observed in this study, this strategy might be universally used against all pathogenic bacteria, including MDR bacteria.

## 3. Conclusion

In conclusion, we report for the first time the ability of pathogenic bacteria to form intracellular MNPs. The bacteria biosynthesized MNPs in presence of iron and zinc precursors *in vitro*. The superparamagnetic nanoparticles composed of zinc ferrites, bestowed the bacteria with magnetic-field induced motility, a property that can be used for bacterial separation. On exposure of the bacteria to AMF, the heating produced by the MNPs induced bacterial cell death. The ability of these elements to enter the biofilms and form MNPs, and the subsequent susceptibility of the bacteria in the biofilms with AMF treatment, suggested the possibility of using the biosynthesized MNPs to overcome the enormous challenges facing biofilm eradication.

Lastly, the identification of MNPs in the bacteria isolated from infected tissues, established the existence of magnetic bacteria in the host. The therapeutic potential of these MNPs to induce magnetic hyperthermia-induced cell death was confirmed using infected tissue specimens. The proposed approach of controlling the virulent bacteria *in vivo* with AMF *per se*, needless to say is simple and safe, with no likelihood of developing resistance, as seen with antibiotics. Moreover, this approach can be utilized to combat co-infections and/or superinfections in patients with Covid-19.

## 4. Experimental Section/Methods

### 4.1 Materials

LB media, zinc gluconate (TCI chemicals Pvt. Ltd, India), ferrous chloride (Sigma-Aldrich) were procured from the respective commercial sources. *Staphylococcus aureus-*ATCC 25923; *Escherichia coli-*ATCC 25922; *Pseudomonas aeruginosa-*ATCC 27853 *and Klebsiella pneumoniae-*ATCC 700603 were purchased from ATCC, USA. Clinical specimens were obtained from PGIMER, Chandigarh after receiving the requisite ethics committee approval.

### 4.2 Bacterial culture with FeCl_2_ and zinc gluconate

A single colony from each plate of the respective Gram-positive and Gram-negative bacteria (*Staphylococcus aureus*; *Escherichia coli; Pseudomonas aeruginosa and Klebsiella pneumoniae-*) was grown in a suspension culture for 2 h in 5 mL of LB. This was transferred to 200 mL of fresh LB medium and grown until OD_550_= 0.5-0.6 (∼5-6 h). For treatment, sterilized salt solutions of 1mM each of FeCl_2_ and zinc gluconate were added to the culture. The resulting suspension was incubated at 37°C with shaking at 180 rpm. At the end of treatment (24-48 h), the bacterial cultures were centrifuged at 1000 x *g*, at 4°C and gently fixed for further analysis. To determine viability, the respective bacterial cultures after 48 h treatment were plated on agar plates to check for growth. Perl’s Prussian blue Staining and Gram Staining were done using standard protocol. ROS levels in the bacteria were determined using carboxy-H_2_DCFDA (Sigma-Aldrich) assay. H_2_O_2_ was used as positive control for ROS induction. Sepsis-positive blood samples were obtained from the Department of Medical Microbiology, PGIMER, Chandigarh, India after receiving approval from the ethics committee (Micro-CC-46).

### 4.3 Magnetotaxis Study

To check the influence of magnetic field on bacterial culture, a drop of the individually treated bacteria was placed on a slide and a neodymium bar magnet was placed on the microscope stage near the drop, with the axis of the magnet parallel to the plane of the slide. The movie of the migration of the microbes towards the magnet was recorded at 15 X speed and 40X magnification under bright field (OLYMPUS, DC73).

### 4.4 Characterization

For TEM analysis, the respective bacterial cells were fixed with 4% formaldehyde solution and drop casted on TEM grids. The grids were negatively stained using Uranyl acetate (.05%), washed and air dried. TEM analysis was carried out at 120 KV (JEOL JEM 2100 transmission electron microscope). For further characterization of nanoparticles the bacterial lysate was used. Nanoparticle size, EDAX, elemental mapping was evaluated using TEM. HR-TEM, SAED pattern and inter planar spacing was used to identify the composition of the nanoparticles. The crystalline structure of nanoparticle was determined by Bruker D8 Advanced X-ray diffraction (XRD) system, using Cu-Ka radiation source from 20 to 80 (2h) with an increment of 0.02 min^-1^ with the respective microbes after calcination at 300 ^0^C. Fourier transform infrared (FT-IR) of lyophilized samples of the treated bacterial culture was recorded by ATR using VERTEX 70 FT-IR (Bruker) from 4000-400 cm^-1^ at 4cm^-1^ resolution. Quantitative estimation of iron and zinc in the bacterial cells (normalized to the number of cells/well) was done using ICP-MS (Agilent Technologies 7700 Series). Magnetic measurements to determine the magnetic properties of the nanoparticles and measurements of temperature dependence of magnetization (ZFC-FC curves) were conducted using a SQUID (Superconducting Quantum Interference Device magnetometer).

### 4.5 ROS determination using carboxy-H_**2**_**DCFDA**

Treated as well as untreated bacterial cells of *S. aureus* and *E. coli were* washed with phosphate-buffered saline (PBS) and treated with Carboxy-H_2_DCFDA dye at a final concentration of 50 µg/ml in PBS. The cultures were incubated in dark for 30 minutes. For positive control, H_2_O_2_ (200µM) was used. ROS levels were assessed by the detection of green fluorescence (excitation filter = 482 ± 35 nm) using fluorescence plate reader. All DCFDA fluorescence intensities were calculated relative to H_2_O_2_ control.

### 4.6 Response to hyperthermia

The magnetic hyperthermia efficiency of the biosynthesized nanoparticles was tested using the D5 heating system (nB nanoscale Biomagnetics, Zaragoza, Spain). The bacterial cells, *E. coli* and *S. aureus* after the treatment with FeCl_2_ and Zn gluconate were then placed in DM2 system and subjected to an alternating current (AC) magnetic field (H) f=164, 347 kHz with field amplitude of, 400 Oe to determine the rise in temperature with a constant target-temperature program feedback at 43°C set point for 30 min. A fluoro-optic thermometer fiber probe was used to probe the temperature every 0.2 s after switching on the magnetic field. The difference in temperature (Δt) was noted as the temperature change observed between treated and untreated cells. The experiments were run in triplicates and each run time was set to 30 min. In order to calculate the reduction in cell number, *E. coli* and *S. aureus* post-AMF treatment were plated on LB agar plates (dilution factor 10^4^). After 24 h of incubation, the colonies were counted and expressed as the number of cells versus AMF treatment.

### 4.7 Effect on Biofilms

*S. aureus* and *E. coli* cells were cultured till they reached an OD_600_ of 0.6 was obtained, which corresponds to 3×10^8^cells/ml. A dilution in nutrient media was made and 1ml of cell suspension was plated on glass coverslips and incubated at 37°C for 72 h. After incubation period was over and a layer of biofilm was formed, Fe and Zn solution were added to the cells and further incubated for 36h. Cells were subjected to hyperthermia at 405 KHz, 400 Oe for 30 min. In order to check the response of AMF treatment on cells, biofilms were subjected to live/dead staining after incubation with 5µl (5mg/ml) stock of FDA (Fluorescein diacetate, used for staining live cells green) and 2.5µl (2mg/ml) of PI (propidium iodide/used for damaged cells red) and visualized using Olympus microscope (OLYMPUS, DC73).

### 4.8 Bacterial susceptibility to antibiotics

The susceptibility of the clinical specimens to multiple antibiotics was assessed as per the latest Clinical and Laboratory Standards Institute (CLSI), To compare the reduction in bacterial viability, the respective bacteria were grown to the mid-log phase and then diluted to a final concentration of 5 × 10^5^ CFU/mL. The cultures were treated with ciprofloacin/vancomycin (2 μg /mL). After O/N incubation at 37°C, the turbidity at OD_600_ was measured by the microplate reader. A 10-fold serial dilutions of the aliquot were plated onto Müller-Hinton agar in triplicate. The bacterial colonies (CFU) were counted after 24 h incubation at 37 °C.

### 4.9 Statistical Analysis

Each statistical data is derived from a sample size of three, and the data is presented as mean ± standard deviation (SD). Graph Pad Prism 6 was used to analyze data and declared statistically significant if P ≤ 0.05 as determined by one-way or two-way ANOVA test keeping a 95% confidence interval.

## Contributions

S.K. Visualized and compiled the manuscript. J.T., P.M., N.S., R.S. and H.S.R. Supported in the characterization, ROS and hyperthermia studies. V.C. conducted the biofilm studies. V.P Analyzed the characterization data; R. K. & V.G supported analysis of clinical microbial cultures, investigations and manuscript editing. D.G. Conceptualized, supervised, secured funding and edited the manuscript. All authors read and approved the final manuscript.

## Acknowledgements

The authors thank Dr. Chayan K. Nandi (IIT, Mandi), Dr. Suvankar Chakraverty, INST and Dr. Nitin Singhal, NABI for their support. Ms. Gurleen Kaur is acknowledged for assistance with graphics. The authors thank Department of Biotechnology (BT/PR22067/NNT/28/1163/2016), Department of Science and Technology (SERB/F/755/2019-2020), India for partially funding the project.

## Corresponding author

Correspondence to Deepa Ghosh

---

TOC

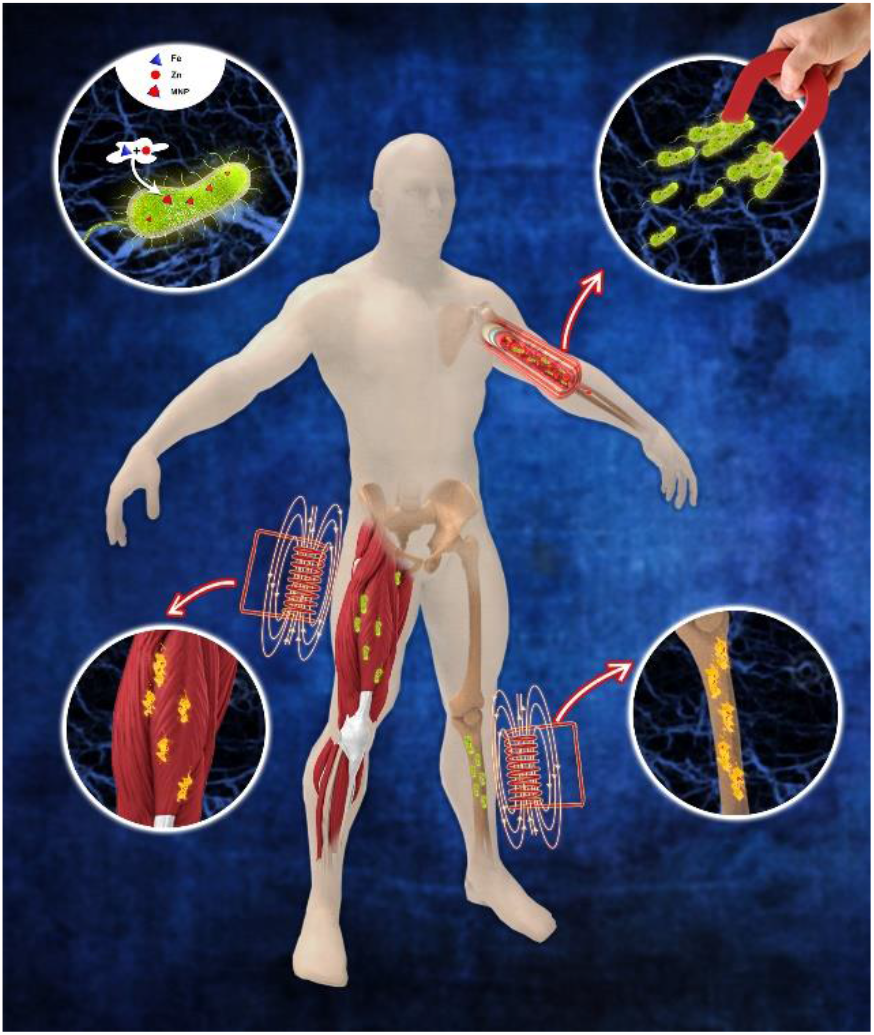

**The formation of intracellular MNPs in pathogenic bacteria and its applications**. Intracellular magnetic nanoparticle (MNPs) are formed in pathogenic bacteria cultured with iron and zinc salts. Bacteria existing *in vivo*, acquire these elements from the host and form MNPs. The MNPs facilitate magnet-induced bacterial motility and hyperthermia-induced cell death on exposure to alternating magnetic field (AMF). Drug-resistant infectious bacteria in tissues/biofilms can be killed with AMF.

## Supporting Information

**Figure S1:**
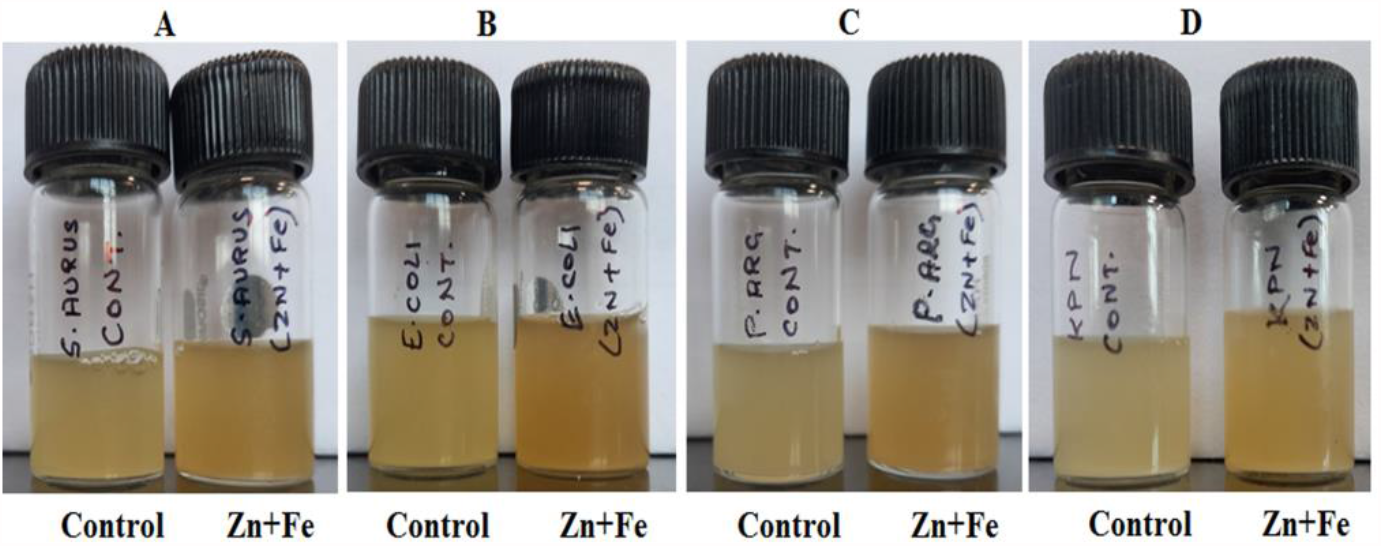
Bacteria treated with/without Fe and Zn. (A) *S. aureus;* (B) *E. coli;* (C) *P. aeruginosa; K. pneumoniae*. Untreated bacteria served as control. Color change observed after 36 h treatment.

**Figure S2:**
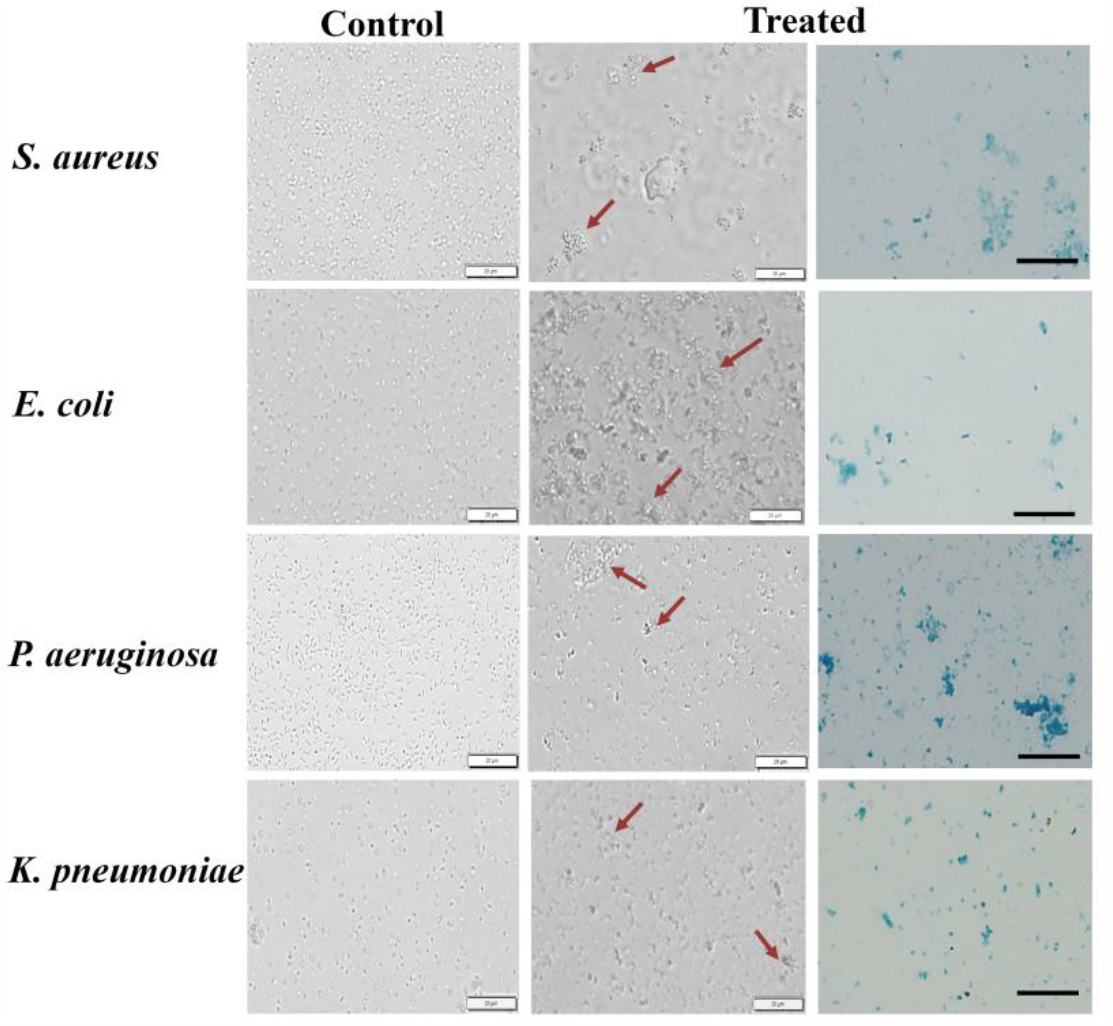
Bacterial response to Fe and Zn. Bright field images of untreated (Left column) and treated bacteria (middle column). Arrows show bacterial aggregates. FeCl2 and zinc gluconate treated bacteria stained with Perl’s Prussian blue (Right column); Scale bar represents 20 µM.

**Figure S3:**
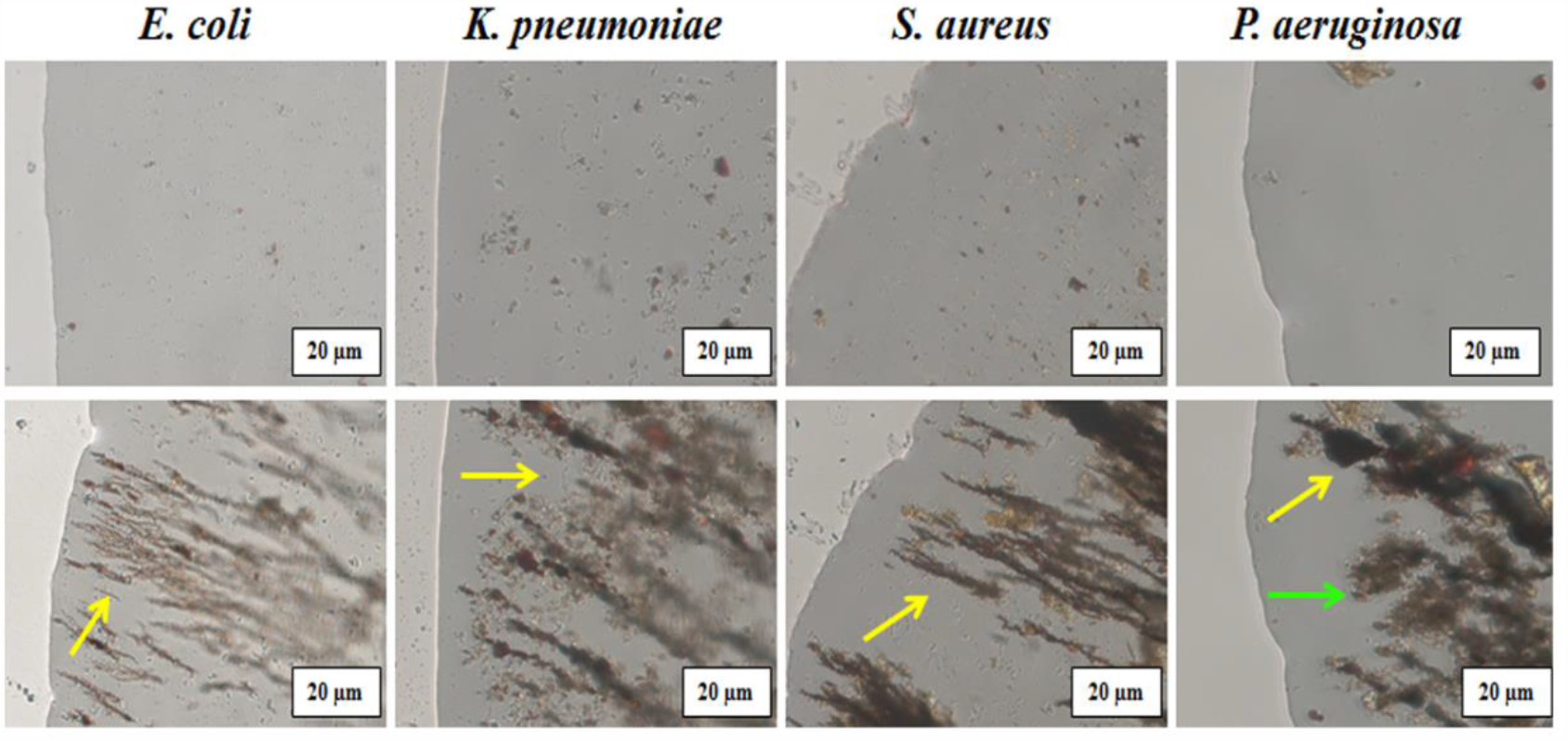
Magnetic-field induced migration of MNPs. Lysates of bacteria treated with Fe & Zn. Upper panel: Microscopic images of the periphery of a drop of lysate. Lower panel: Images of magnet-induced aggregation of nanoparticles. Yellow arrow represents magnetic nanoparticles and green arrow represents cell debris. **M1-M6** **Magnetotaxis Videos** **M1:** Movie of untreated *S. aureus*. **M2:** Movie of *S. aureus* treated with FeCl_2_ + Zn gluconate. **M3:** Movie of untreated *E. coli*. **M4:** Movie of *E. coli* treated with FeCl_2_ + Zn gluconate. **M5:** Movie of untreated *P. aeruginosa*. **M6:** Movie of *P. aeruginosa* treated with FeCl_2_ + Zn gluconate.

**Table ST1:** Estimation of Fe and Zn. IC-PMS analysis of Fe and Zn in the lysates of control and bacteria treated with / without FeCl2, and zinc gluconate. Results represent values obtained from 3.8 x10^8^ CFU

| Name of the Bacteria | Conc. of Fe µg/ml | Conc. Of Zn µg/ml |
| --- | --- | --- |
| *E.coli* (Control) | 0.164±10 | 0.198±2 |
| *E.coli* (FeCl<sub>2</sub>) | 0.179±7 | 0.213±4 |
| *E.coli* (Fe+Zn) | 28.33±3 | 0.602±10 |
| *S.aureus* (Control) | 0.116±12 | 0.227±5 |
| *S.aureus* (FeCl<sub>2</sub>) | 0.169±6 | 0.283±8 |
| *S.aureus* (Fe+Zn) | 12.8±4 | 0.512±7 |

**Figure S4:**
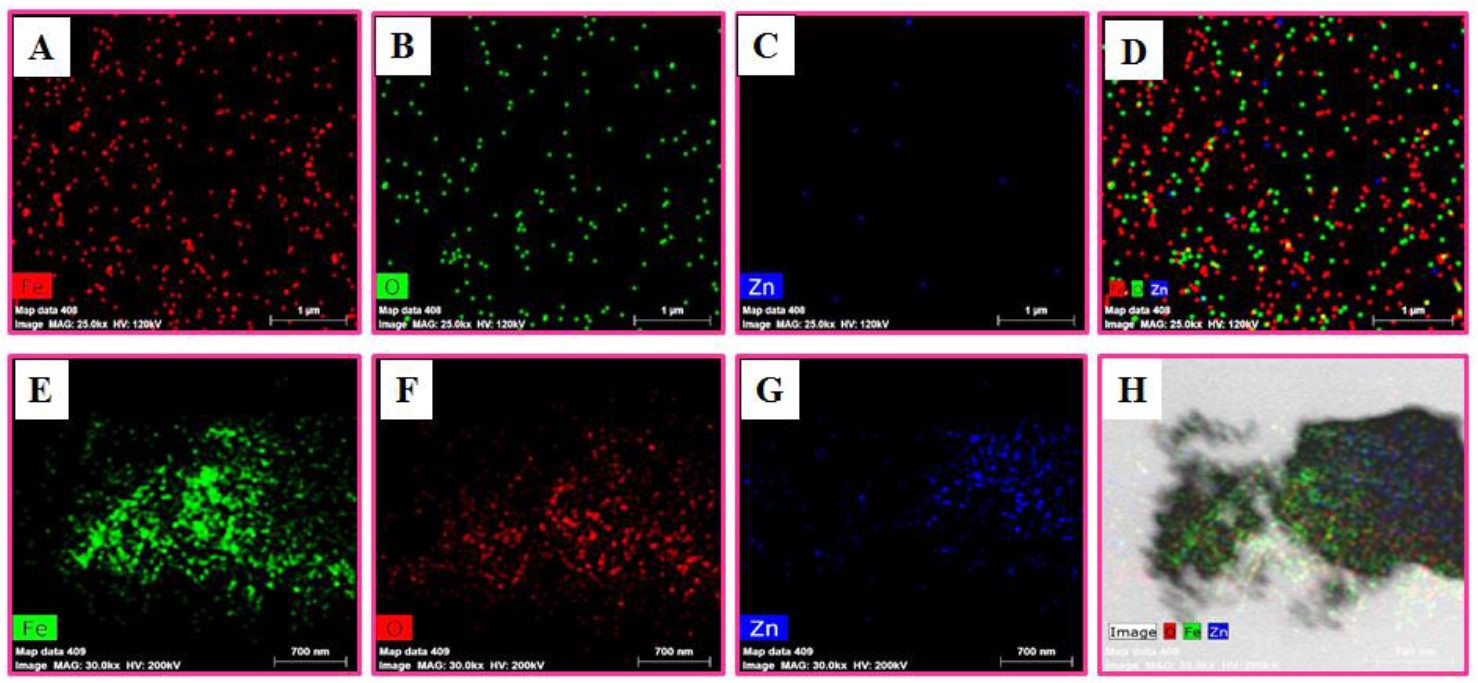
Elemental mapping using TEM. Distribution of iron, oxygen and zinc in treated bacteria. Upper panel represents *S. aureus* (A-C) and Lower panel *E. coli* (E-G). D & H indicate a combination of (A-C) and (E-G) respectively.

**Figure S5:**
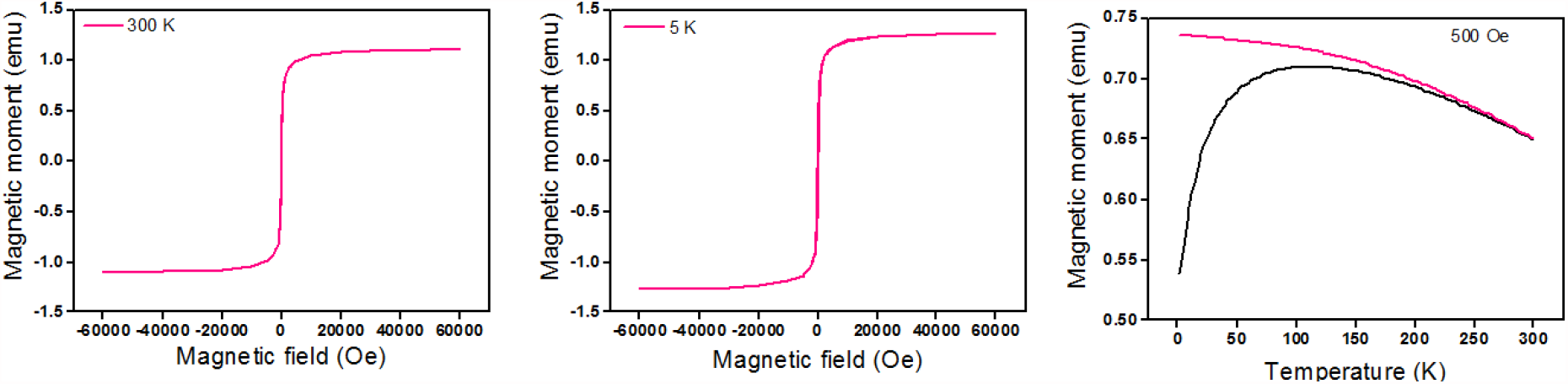
Magnetic measurements of nanoparticles biosynthesized in *E*.*coli*. Magnetization versus magnetic field measured at (A) 300 K, (B) 5K and (C) Measurements of Temperature Dependence of Magnetization (ZFC/FC curves).

**Figure S6:**
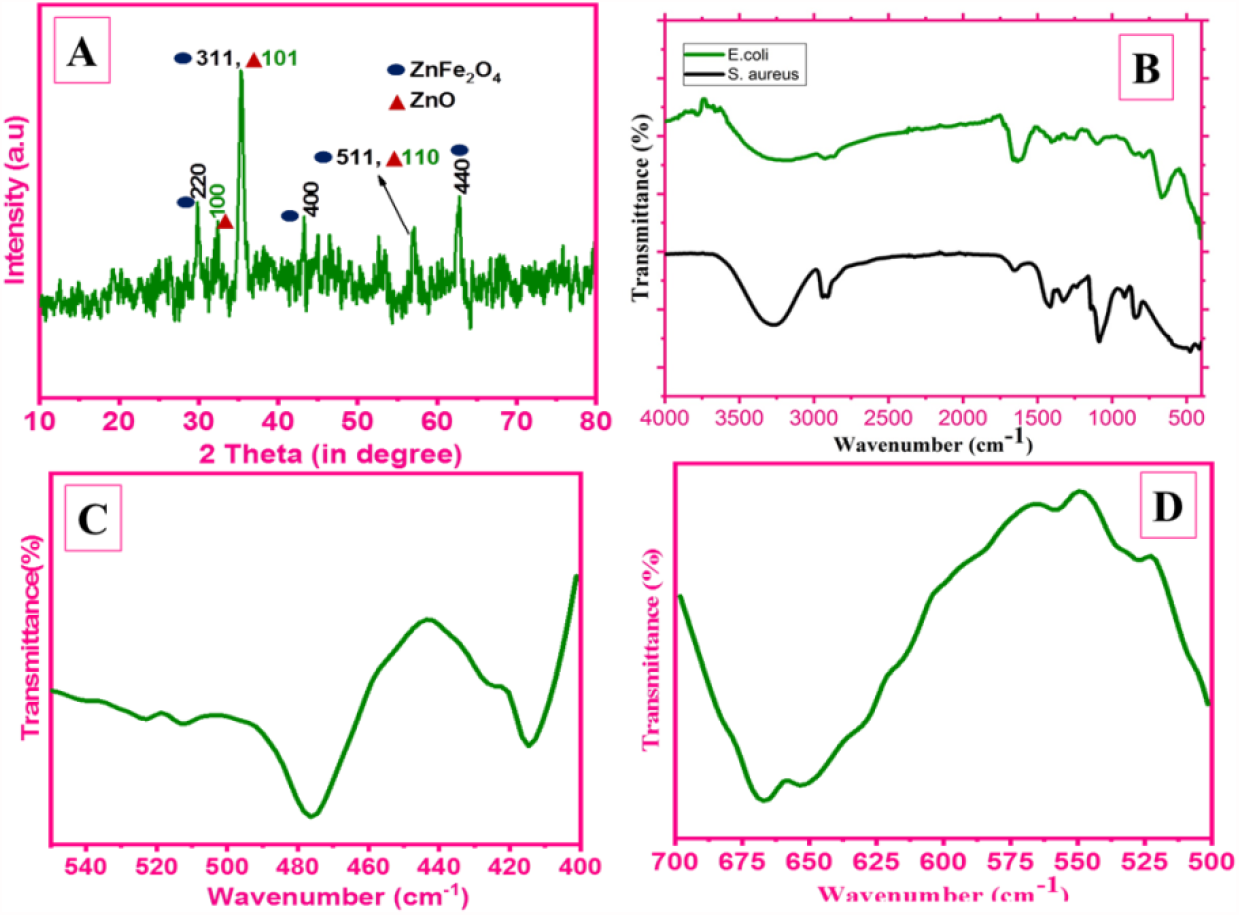
Characterization of Fe and Zn treated bacteria. (A) XRD diffraction pattern of *S. aureus* after calcination. (B) FT-IR of the treated bacteria after lyophilization of *S. aureus* and *E. coli*. (C) Peaks observed in the region from 476-417 cm^-1^ represent Zn-O bond (D).Peaks in the region from 670-550 mode represent Fe–O bonds.

**Table ST2:**
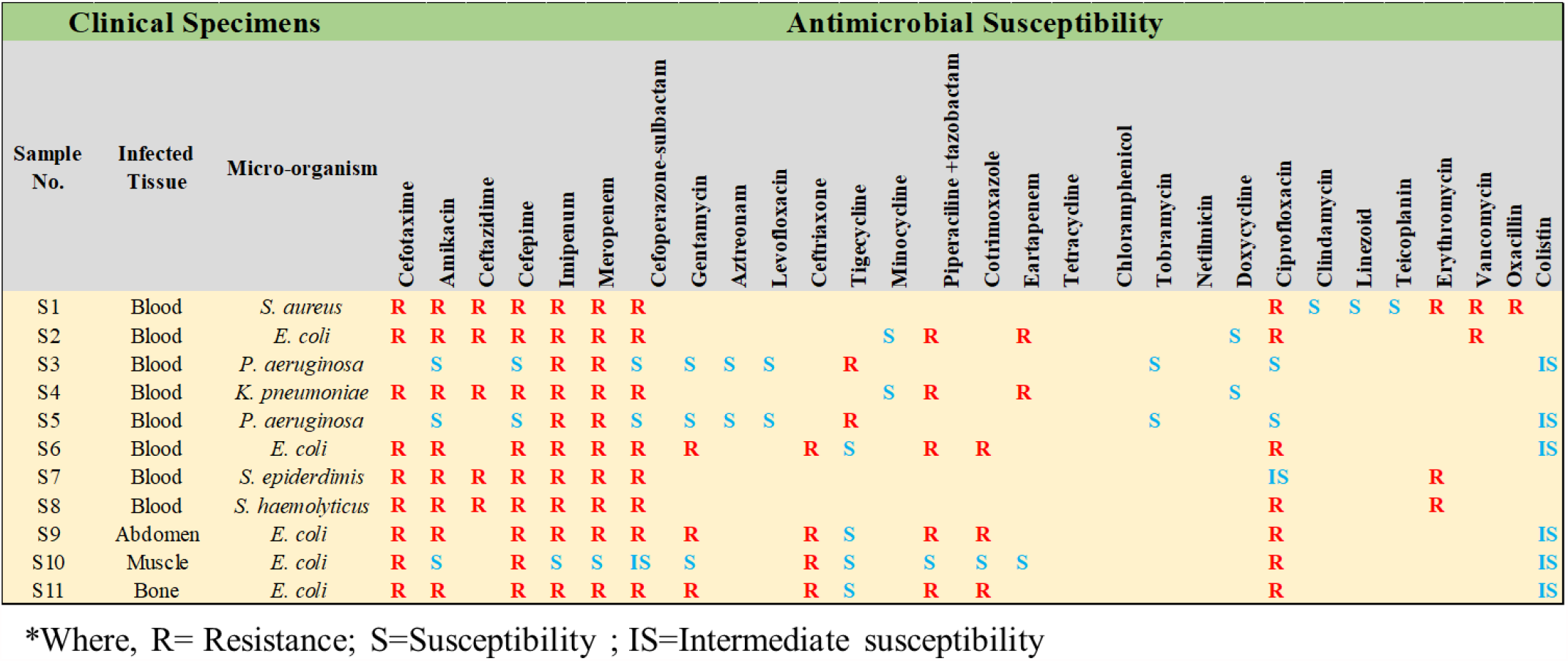
Table represents the various infected specimens analyzed and its susceptibility to antibiotics

**Figure S7:**
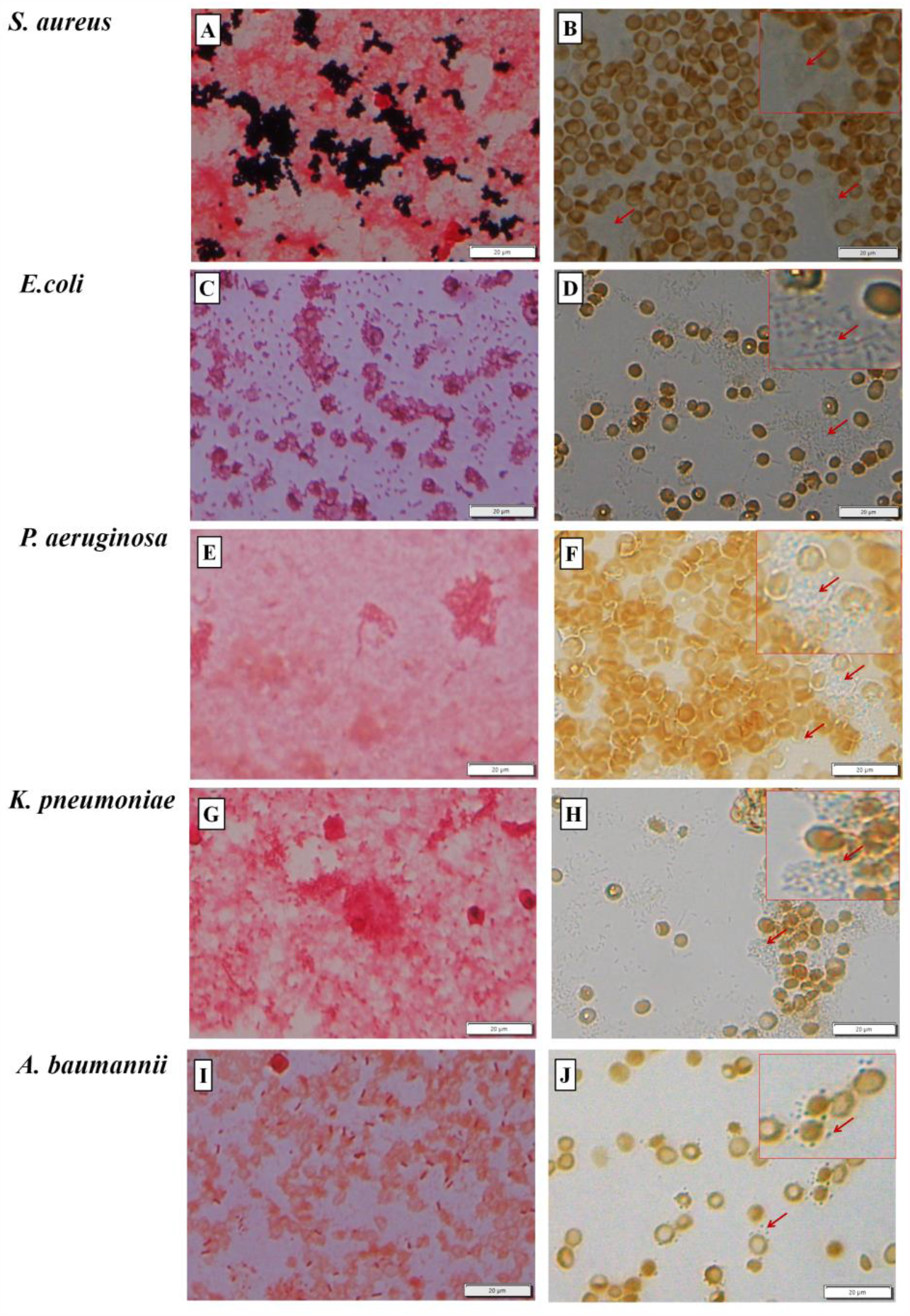
Clinical strains tested for the presence of iron oxide. (**A, C, E, G, I**). Gram-staining of respective bacteria. (**B, D, F, H, J**) Blood smear of respective bacterium stained with Prussian blue. Inset shows the magnified image. Arrows indicate blue color obtained after Prussian blue staining. Scale bar represents 20 µm.

**Figure S8:**
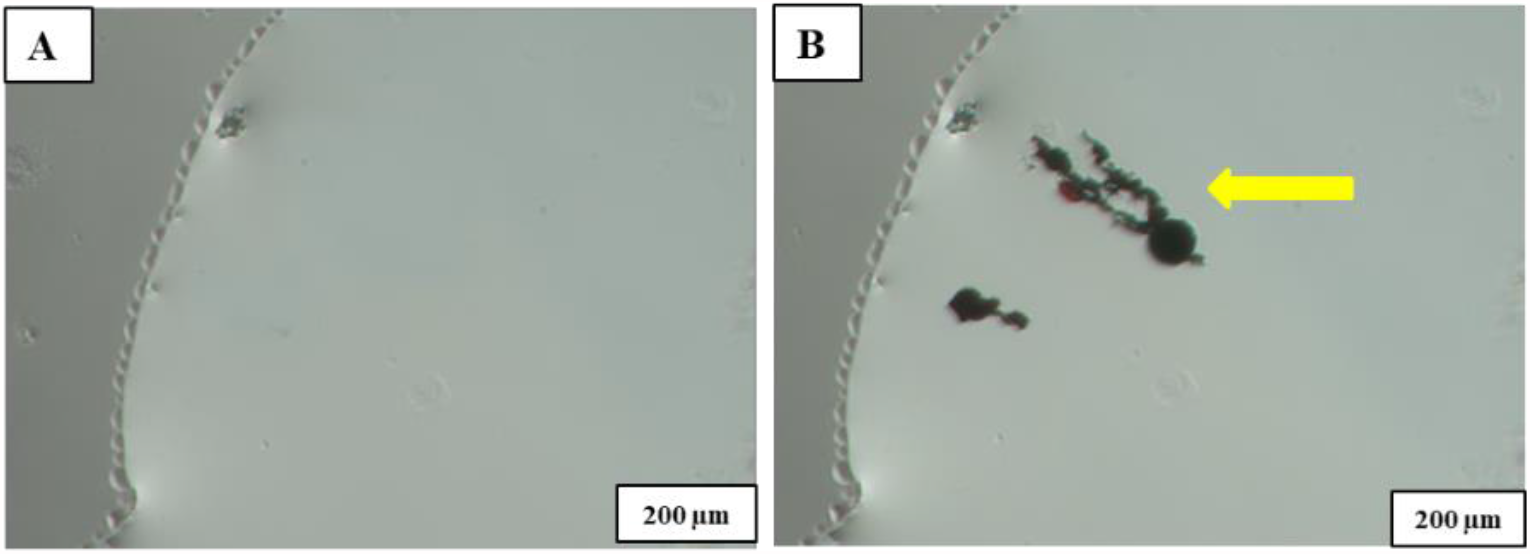
Magnetic-field induced migration of MNPs obtained from *S. epiderdimis*: **A**. Microscopic image of the periphery of a drop of *S. epiderdimis* lysate. **B**. The lysate exposed to magnet. Arrow indicates the alignment of MNPs along the magnetic field lines.

**Figure S9:**
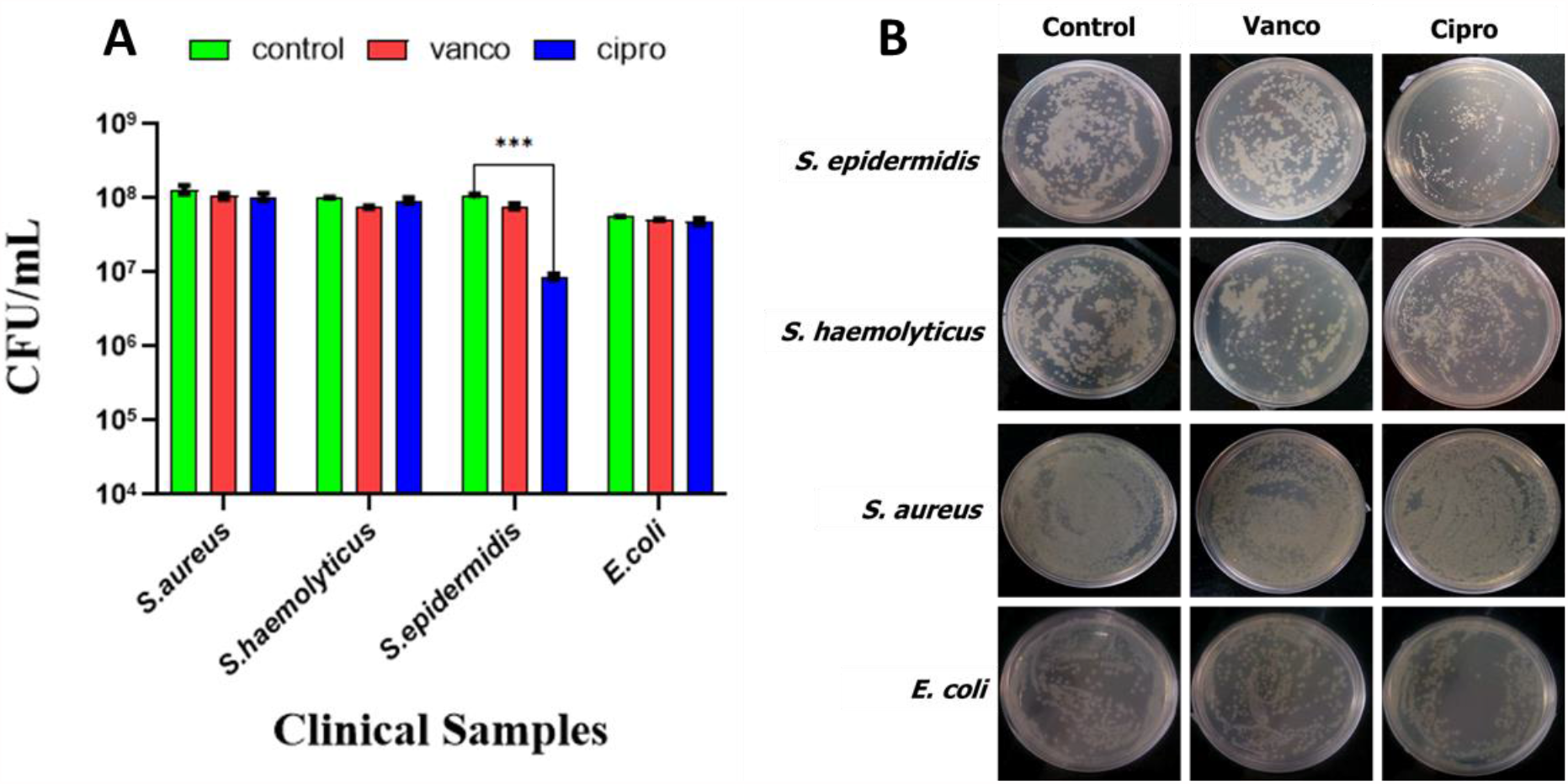
Susceptibility of bacteria (clinical specimens) to vancomycin and ciprofloxacin. The respective bacteria isolated from the infected blood specimens were treated with vancomycin/ ciprofloxacin (2 μg/ml) for 24 h. **(A)** The number of bacterial colonies obtained after respective treatment. The data represents the mean ± SD obtained from 3 experiments in which N=3 in each group. Two-way ANOVA Bonferroni multiple comparison test where *** represents p-value <0.001 **(B)** Representative images of the bacterial colonies grown on agar plates.

**Figure S10:**
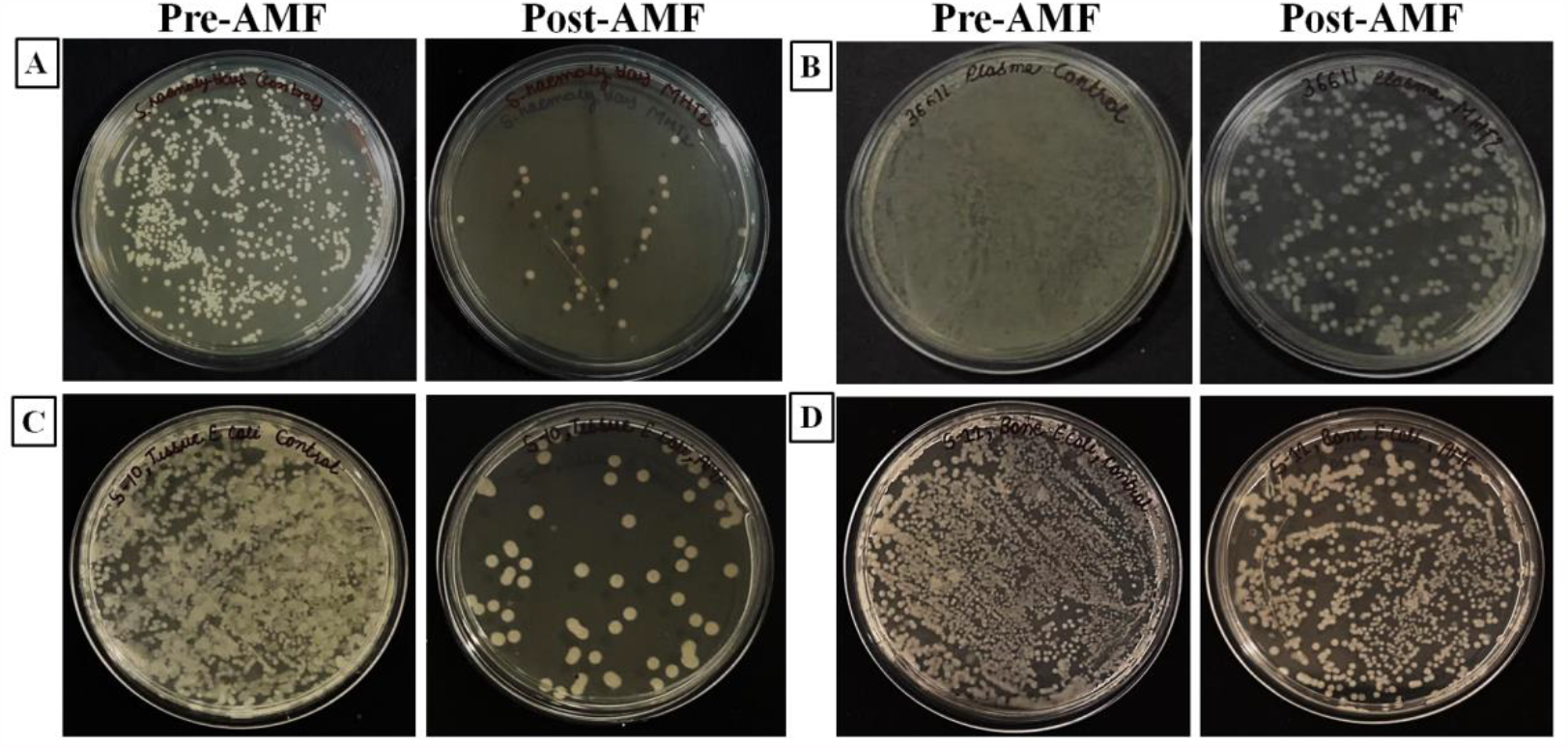
Effect of AMF on infected clinical specimens: Bacterial colonies obtained from pre- and post-AMF exposed samples A **& B** represent *S. haemolyticus* and *E. coli* infected blood samples exposed to AMF (347 KHz, 400 Oe); **C & D** represents *E*.*coli* infected muscle and bone specimens exposed to AMF at 405 KHz, 400 Oe respectively.

